# Gilda: biomedical entity text normalization with machine-learned disambiguation as a service

**DOI:** 10.1101/2021.09.10.459803

**Authors:** Benjamin M. Gyori, Charles Tapley Hoyt, Albert Steppi

## Abstract

**Summary:** Gilda is a software tool and web service which implements a scored string matching algorithm for names and synonyms across entries in biomedical ontologies covering genes, proteins (and their families and complexes), small molecules, biological processes and diseases. Gilda integrates machine-learned disambiguation models to choose between ambiguous strings given relevant surrounding text as context, and supports species-prioritization in case of ambiguity.

**Availability:** The Gilda web service is available at http://grounding.indra.bio with source code, documentation and tutorials are available via https://github.com/indralab/gilda.

## 1 Introduction

Named entity recognition (NER) and named entity normalization (NEN) are central tasks in extracting knowledge about genes, proteins, metabolites, small molecules, biological processes, and other entities of interest from unstructured text in the biomedical literature. The role of NER is to identify an entity string (e.g., “JNK-1”) in text, and NEN, or *grounding*, is to choose an appropriate namespace and identifier corresponding to an entity string (e.g., HGNC:6881 for “JNK-1”).

While UniProt, ChEBI, GO, and other resources make available curated lists of names and synonyms for different entity types, exact searches over their thesauri have been shown to provide insufficient coverage due to the diversity of biomedical nomenclature [1]. It is further necessary to account for variability in spaces, dashes, capitalization and the expansion of certain characters (e.g., Greek letters) with respect to these lexicalizations to find relevant matches. A further challenge is that of ambiguity, namely, that several entities can share the same name or synonym, and it is only based on broader context (a surrounding sentence or paragraph) that one can resolve which sense is implied in the given setting.

While NEN is typically integrated in systems solving more complex tasks such as full-text annotation [2] or relation extraction [3], there are many ways in which NEN is useful as a standalone tool, including in the context of interactive search interfaces (finding identifiers for search terms entered by users), data analysis (finding identifiers for data entries, e.g., drugs), and modeling (assigning identifiers for modeled concepts).

Here, we present Gilda, a software tool and web service which integrates multiple biomedical lexical resources and implements a scored string matching algorithm (parts of which were adapted from the text tagger in [4]) to names and synonyms across entries in these resources. Importantly, Gilda makes available over 1,000 machine-learned disambiguation models for strings representing multiple ambiguous entities, and can apply these models as part of the scoring process given surrounding text as context. Gilda achieves state of the art performance on several of the BioCreative VI NEN benchmarking tasks and is competitive on the rest.

Gilda is available as a public REST web-service at http://grounding.indra.bio, and as an open-source Python package under a BSD license at https://github.com/indralab/gilda.

## 2 Results

### Lexical Resources

Gilda integrates over 1.8 million names and synonyms from 9 ontologies and lexical resources for the purpose of name matching (Supplementary Table 1). In addition to standard resources such as UniProt, Gilda integrates FamPlex, a resource providing lexicalizations for protein families and complexes (shown to be a key missing component in named entity normalization systems) as well as over two thousand lexicalizations for entities that were manually curated using a frequency-ranked list of strings mined from the literature that did not appear in other standard resources [1].

Each of the resources listed in Supplementary Table 1 are processed to extract a list of *terms*, with each term carrying the following information, illustrated via the example of “MEK1”: (i) Namespace (e.g., HGNC); (ii) Identifier within the namespace (e.g., 6840); (iii) Text name (e.g, MEK1); (iv) Type of text name (e.g., synonym); (v) Canonicalized text name (e.g., mek1); (vi) Standard name (e.g., MAP2K1); (vii) Source (e.g., hgnc).

To allow extensions and customizations, the Gilda Python package supports instantiation with custom grounding resources, and is able to load terms from ontologies in OWL, OBO, and other standard formats using the PyOBO^1^ package.

### Efficient approximate scored string matching

Gilda implements a grounding algorithm inspired by [4] that allows for efficient approximate matches to any of the terms appearing in the integrated resource table. For a given entity string, the grounding process first generates relevant variants (e.g., by spelling out Greek letters) and then identifies possible matches for each variant. This involves canonicalizing the variant string, and then searching the resource table for the same canonicalized text name. This lookup is efficient since the service loads the resource table in memory, indexed by canonicalized text names in a hash map. A string comparison algorithm then compares the original (i.e., not yet canonicalized) entity string with each matched entry to assign a score. The string comparison algorithm takes the following into account when scoring each match: (i) dashes, hyphens and spaces; (ii) capitalization; (iii) whether the entity is possibly a plural form; (iv) the status of the term that was matched (standard name, synonym, withdrawn entry, etc.). The final score is between 0 and 1, with 1 corresponding to an exact match of a standard name. Further details of the matching and scoring procedure are described in Supplement section 3.2.

### Machine-learned models for context-aware disambiguation

Many of the entries integrated in Gilda’s resource table share the same text name, including over one thousand gene synonyms that are shared across multiple human genes. These entries are textually equivalent and their status (i.e., synonym) is identical, therefore the scores assigned to them for a given input entity string will be the same.

To resolve this ambiguity, we used labeled articles from the biomedical literature to learn models of the context surrounding each member of a set of ambiguous terms that share a synonym (see Supplement section 3.3 for details). Gilda currently contains such disambiguation models for 1,008 ambiguous strings (e.g., “HK4”, “p42”). Gilda also integrates 153 disambiguation models made available by the Adeft system [5]. Adeft models are classifiers that can choose between senses of the most commonly occurring acronyms in biology literature with multiple senses (e.g., “ER”) given surrounding text context.

When Gilda is called with an ambiguous string that one of the models is applicable to, it uses context (for instance, a sentence or a paragraph) supplied by the user to make a prediction and adjust the scores of returned matches. More information about these models can be found in Supplementary section 3.3.

### Cross-species protein disambiguation

In addition to text as context, Gilda also takes an optional species prioritization list as input. The list can be provided directly by the user to express prior knowledge or preference for normalizing to a specific species such as human or yeast. Alternatively, in case the text to be normalized comes from a PubMed-indexed publication, Medical Subject Headings annotations associated with the publication can be used to automatically derive a species preference order.

### Benchmarks

We benchmarked Gilda on a grounding task with the BioCreative VI Track 1 (Bio-ID) data set from [6]^2^. When used as a Python package, Gilda was able to ground over 20 thousand strings per second in this benchmark (Supplementary Table 4). Gilda achieved state of the art *F*_1_ for proteins (.693 for human .616 for non-human vs. .445 from [7]), cellular components (.504 vs. .476 from [8]), and small molecules (.620 vs. .591 from [8]). It underperformed for species (.586 vs. an average .623 over several configurations from [9]) cells/cell lines (.595 vs. .740 from [8]) and tissues (.446 vs. .633 from [8]) likely due to gaps in the lexical resources in Gilda covering those entity types. While most previous works generated static entity-type-specific models, Gilda uses a single general model for all entity types which can be readily updated on new releases of its underlying lexical resources and extended to new tasks by incorporating additional lexical resources. However, we acknowledge that its generalization has a negative impact on Gilda’s precision in comparison to entity-type-specific models.

We also benchmarked Gilda on the evaluation data set described in [1] where a random sample of 300 named entities extracted from a subset of literature were curated manually and compared to automated grounding. We found that using Gilda improved precision from .900 to .943, and recall from .850 to .895 compared to the results reported in [1] (see also Supplementary Table 3).

### Availability and integrations

Gilda is available as a Python package through PyPI, a Docker image through DockerHub, and as a web application with a RESTful API at https://grounding.indra.bio. Gilda is integrated into INDRA [10] to ground entity texts from knowledge sources where grounding is missing and to disambiguate entity strings from text mining. Gilda is also integrated into multiple web services and a human-machine dialogue system (see Supplement section 3.7).

## 3 Supplementary Material

### 3.1 Lexical Resources

Gilda (version 0.6.1) integrates terms extracted from the resources listed in Table 1. Gilda can also be instantiated with other custom resources loaded as Terms.

**Table 1.** Resources integrated in Gilda.

| Resource | Entity types | # strings | Reference |
| --- | --- | --- | --- |
| HGNC | Genes | 105,109 | [11] |
| UniProt | Proteins | 437,645 | [12] |
| ChEBI | Small molecules, metabolites | 321,599 | [13] |
| GO | Biological processes, complexes, cellular locations | 143,309 | [14] |
| MeSH | Multiple entity types | 756,326 | [15] |
| FamPlex | Protein families and complexes | 3,051 | [1] |
| EFO | Cell types, anatomy, disease, etc. | 30,507 | [16] |
| HPO | Phenotypic abnormalities | 30,201 | [17] |
| DO | Diseases | 23,793 | [18] |

### 3.2 String Matching and Scoring

Keys into Gilda’s grounding terms are stored in a canonical form. Given a string *Y*, the canonical form 𝒞 (*Y*) is obtained by replacing all contiguous runs of white-space characters with a single ASCII space, converting all text characters to lower case and removing all ASCII and Unicode dash characters.

Given a string *X* containing the raw text of an entity that is to be grounded, Gilda generates a set of lookup strings ℒ(*X*) and searches for them among the canonicalized keys in the lexicon. One lookup string in ℒ(*X*) is the canonical form 𝒞 (*X*), but an additional set of lookup strings are included to allow for more flexible matching. Given *X*, lookup strings are generated for the canonical forms of each of the following strings.

- *X*.
- *X* after all ASCII and Unicode dashes have been replaced with spaces (example: “EGF-receptor” -¿ “EGF receptor”).
- *X* after all spelled out Greek letters have been replaced with their single character Unicode equivalents.(example: “PKC-alpha” -¿ “PKCA”)
- *X* after all spelled out Greek letters from a subset with close Latin equivalents have been replaced with the single character Latin equivalent. (example: “IKK-*β*” -¿ “IKKB”)
- *X* after all Unicode Greek characters have been replaced with the spelled out Latin form. (example: “GSK3-*β* -¿ “GSK3-beta”)
- The depluralized form of *X*, if *X* is determined to be in a possible plural form through rule-based pattern matching. (example: “RAFs” -¿ “RAF”)

Matches are then scored based on the “status” of the matching entry in the lexicon and based on a string comparison score computed between the original uncanonicalized agent text *X* and the original uncanonicalized entry *Y* in the lexicon corresponding to the match.

The possible statuses for entries in the lexicon are listed below. Each status is assigned a numerical score encoding its priority.

- **Assertion:** A name or synonym for an entity which has been manually curated as unambiguous. *Score 4*
- **Name:** The standard name for an entity within the given ontology or lexical resource. *Score 3*
- **Synonym:** A known synonym for an entity that is listed within the given ontology or lexical resource. *Score 2*
- **Previous:** A term which was previously the standard name for an entity in a given ontology or lexical resource. *Score 1*

Given a raw agent text *X* and the associated raw agent text *Y* for a matching entry, the string comparison score 𝒮_string_ (*X, Y*), is a numerical score between 0 and 1. If the entry corresponding to *Y* has status score 𝒮status, and disambiguation score then the overall score for the match is given by

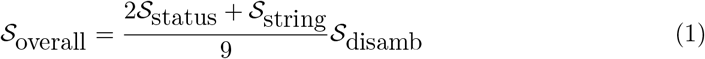

which yields a numerical score between 0 and 1. Here disamb is the probability assigned by a disambiguation model if one was available and applied, or 1 (i.e., no change to the overall score) if disambiguation was not applied. Scoring of individual matches, i.e., the calculation of 𝒮_string_, follows the text tagging algorithm in [4] and is described in detail at http://trips.ihmc.us/TextTagger/docs/README.xhtml#sec-5.7 (see formula for “case-dash-score”). To summarize the key details here: the score is calculated as a normalized linear combination of six sub-score terms. Five of the sub-score terms penalize different patterns of capitalization mismatches between the two strings, while the sixth term penalizes dash-mismatches between the strings.

### 3.3 Disambiguation Models

It is common for there to be multiple distinct groundings for the same entity text within Gilda’s resources. One example of such an ambiguity is the overlapping gene synonym “DAP4” for both DLGAP4 (DLG associated protein 4) and THAP12 (Death associated protein 4). As of Gilda v0.6.1, there are 3,801 synonyms that are shared between at least two distinct human genes and 573 synonyms that are shared between 3 or more distinct human genes.

Gilda offers the feature to disambiguate between the different senses of an entity text through a set of trained logistic regression models, using context provided by the user. Gilda v0.6.1 provides models for disambiguating 1,008 different entity texts and also makes use of models for 172 entity texts provided by the Adeft package [5], allowing for disambiguation of a total of 1,180 ambiguous entity texts. Below we describe the process used to train Gilda’s models and select them for inclusion.

For Gilda, disambiguation is formulated as a classification problem. Gilda adopts the approach of training one classification model per ambiguous agent text and makes the one sense per discourse assumption [19] that an agent text is unlikely to be used with multiple senses within the same document. Within this approach to disambiguation, the key challenge is, for each ambiguous agent text, to produce a corpus of documents where the agent text appears and where each document is labeled with the sense in which the agent text is being used in that document. Producing such corpora of documents can be challenging and is often a bottleneck. Gilda leverages existing annotations of documents from the Entrez Gene database [20] and MeSH [15] by relaxing the requirement that the document uses the given agent text, only requiring that it has been annotated as being relevant to one of the senses of interest.

We identified 1,287 ambiguous agent texts with senses for which a sufficient number of labeled documents were available in Entrez and MeSH. Agent texts with at least one ChEBI grounding were excluded, due to difficulties producing labeled documents for chemicals and because chemical and gene synonyms often overlap. Using Adeft’s Scikit-learn [21] wrapper with default parameters, we trained logistic regression models for each of these agent texts using term frequency-inverse document frequency (tf-idf) [22] vectorized unigrams and bigrams as features, performing 5-fold cross-validation in order to estimate generalization error. We evaluated model performance with Scikit-learn’s macro-averaged *F*_1_ score. For each model, we considered the mean of the macro-averaged *F*_1_ scores from each cross-validation split. A histogram of these *F*_1_ scores over all models can be seen in Figure 1. Reasons for poor performance in some cases include insufficient amounts of training data, and highly similar senses that are difficult to distinguish. We chose a cutoff of 0.7 for macro-averaged *F*_1_ score; all models with score below this cutoff were excluded from deployment.

**Figure 1.**
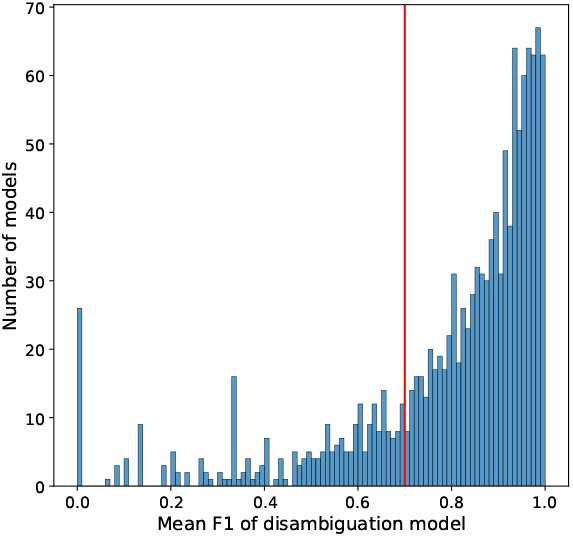
A histogram of *F*_1_ cross-validation scores for all of the trained disambiguation models. A red vertical line indicates the cutoff of 0.7 that was chosen for model inclusion.

### 3.4 BioCreative VI Benchmark Results

We benchmarked Gilda on the BioCreative VI Bio-ID corpus which is available at https://biocreative.bioinformatics.udel.edu/media/store/files/2017/BioIDtraining_2.tar.gz. This corpus consists of 102, 717 entity texts chosen from 570 publications with a curated grounding provided for each entity text (in the “obj” column of the table). Some of the provided groundings, however, do not refer to a specific identifier in an ontology. For example, “protein:GST” appears as an entry in the “obj” column, but it does not represent a specific grounding in an ontology. We classify such groundings as unknown and filter them out to retain only rows that refer to entries in specific namespaces such as NCBI gene, UniProt, GO, Uberon, etc., resulting in a total of 86, 574 rows. The namespaces used in the Bio-ID corpus do not align exactly with ones Gilda uses by default. For instance, Bio-ID provides gene and protein groundings to NCBI gene and UniProt, whereas Gilda uses HGNC for human genes/proteins and UniProt for non-human genes/proteins. Similarly, Bio-ID provides groundings for tissues in Uberon whereas Gilda uses MeSH to ground tissues. We therefore applied equivalence mappings between these namespaces to allow matches to equivalent entries in different ontologies from the one used for a given row in Bio-ID. For example, one row of Bio-ID contains “brain” as an entity text and provides UBERON:0000955 as its grounding; using equivalence mappings between Uberon and MeSH, we also allow MESH:D001921 as an acceptable grounding.

Table 2 shows the results of this benchmark. We calculated precision and recall under two conditions: a strict one in which we only accept a grounding if it is the “top” match returned by Gilda, and a more permissive one in which we score a grounding as correct if “any” of the matches returned by Gilda corresponds to the expected grounding.

**Table 2.** Precision, recall, and *F*_1_ values for Gilda performance on the Bio-ID corpus by entity type. Values are given both for the case where Gilda’s result is considered correct only if the “top” grounding matches and the case where Gilda’s result is considered correct if “any” of its returned groundings match.

| Entity Type | Prec. (top) | Prec. (any) | Rec. (top) | Rec. (any) | $F_1$ (top) | $F_1$ (any) |
| --- | --- | --- | --- | --- | --- | --- |
| Cell types/Cell lines | .695 | .722 | .520 | .540 | .595 | .618 |
| Cellular Component | .587 | .615 | .442 | .463 | .504 | .528 |
| Human Gene | .710 | .795 | .677 | .758 | .693 | .776 |
| Non-human Gene | .655 | .703 | .582 | .624 | .616 | .661 |
| Small Molecule | .683 | .760 | .568 | .632 | .620 | .690 |
| Taxon | .676 | .707 | .517 | .540 | .586 | .612 |
| Tissue/Organ | .575 | .575 | .364 | .364 | .446 | .446 |
| miRNA | .358 | .358 | .282 | .282 | .315 | .315 |
| Overall | .669 | .722 | .561 | .606 | .610 | .659 |

We noted a relatively large gap for human genes between the “top” and “any” conditions and found that there were two factors affecting these scores. First, we found that the Bio-ID corpus often assigns a specific protein grounding to entity texts which are ambiguous, and could refer to a protein family containing the specific protein. For example, “VEGF” in Bio-ID is annotated as the specific gene VEGFA (HGNC:12680) whereas Gilda grounds it to the VEGF protein family in the FamPlex namespace, an entry which represents all human VEGFs. We found that when we accepted matches by Gilda to a FamPlex family or complex of which the Bio-ID-annotated specific protein is a member, *F*_1_-score for human genes was .762, significantly higher than the .669 reported in Table 2 where only a strict match is allowed. Second, we found that the Bio-ID corpus differentiates human and non-human forms of the same gene within the same paper. In several cases, Gilda found the expected species-specific gene but lower in the priority order (which in Gilda’s case is constructed at the paper level).

### 3.5 FamPlex Benchmark Results

We also evaluated Gilda on a manually curated set of 300 entity texts randomly selected from relation extractions by the Reach system [3] on a corpus of 270,000 papers defined by [1]. The corpus consists of abstracts from PubMed and (where available), full text content from PubMed Central, or via the Elsevier text and data mining API covering articles specifically relevant for signaling pathways (for more details, see Table 1 of [1]). We used the grounding results described by [1] as reference (“Reference” row of Table 3) and calculated precision, recall, and *F*_1_ scores, and compared it with groundings produced by Gilda for the same entity strings (“Gilda” row of Table 3). Corresponding to the grounding logic applied in the reference data, in this setting, we ran Gilda without species-disambiguation. We found that using Gilda resulted in a relative improvement compared to the reference both in terms of precision and recall, by 4.8% and 5.3%, respectively.

**Table 3.** FamPlex benchmarking results

| Trial | Precision | Recall | $F_1$ |
| --- | --- | --- | --- |
| Reference [1] | .900 | .850 | .874 |
| Gilda | .944 | .895 | .919 |
| Absolute Improvement | .044 | .045 | .045 |
| Percentage Improvement | 4.8% | 5.3% | 5.1% |

### 3.6 Gilda responsiveness benchmark

We benchmarked the responsiveness (i.e., speed) of Gilda in three settings: (i) when run as a Python package; (ii) when run as a local web service; and (iii) when using the publicly available web service instance. We used all entity texts from the BioCreative VI Bio-ID corpus (the same corpus used for benchmarking in Supplementary subsection 3.4) and measured the average number of groundings performed per unit time with and without context added (i.e., additional surrounding text provided along with the entity text for disambiguation purposes). For benchmarking Gilda in the first two scenarios (when running as a local Python package or a local web service) we used a desktop PC with a 2.5GHz Intel i9-11900 processor and 64 GB of RAM running Ubuntu 20.04 and Python 3.8.10. Results are shown in Table 4.

**Table 4.** Summary statistics on the Gilda responsiveness benchmark on the Bio-ID corpus under three scenarios (local Python package, local web app, and public web app), and with and without context-based disambiguation. The mean requests served per second and its standard deviation are shown.

| Scenario | Context | Requests per Second |
| --- | --- | --- |
| Python package | False | 22050.9 ± 10168.9 |
| | True | 21558.4 ± 10260.3 |
| Local web app | False | 683.0 ± 64.3 |
| | True | 538.7 ± 69.4 |
| Public web app | False | 28.2 ± 0.9 |
| | True | 15.1 ± 1.7 |

While usage as a local Python package was by far the fastest (over 20 thousand entity strings grounded per second), this usage mode may not be ideal in settings where startup time, memory usage and Python dependencies are undesirable. In such cases using the web application is recommended. The local web application performed better than the remote application likely due to the lack of overhead from network communication.

### 3.7 Integrations

Gilda is used in the INDRA Database web application (https://db.indra.bio) which allows searching for statements assembled by the INDRA system [10]. Here, Gilda grounding allows users to enter entity texts in non-standard form (e.g., using informal synonyms) and select an appropriate grounding used for the search, as demonstrated in Figure 2. Gilda is also integrated into the EMMAA dashboard (https://emmaa.indra.bio) where, similar to its integration with the INDRA Database, it supports queries based on user-entered entity texts.

**Figure 2.**
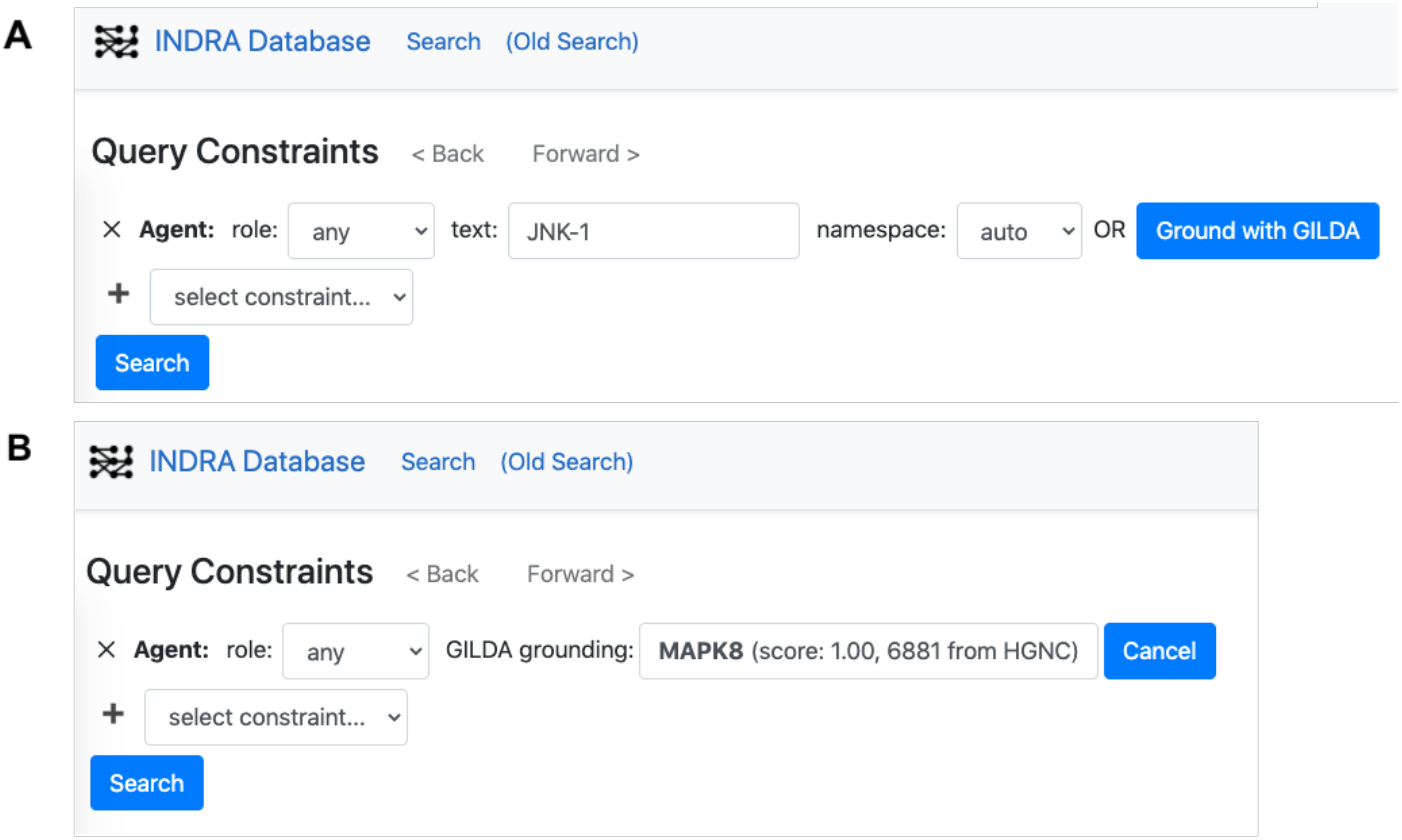
Gilda is used to ground entity texts before submitting search queries to the INDRA Database at https://db.indra.bio. (A) The user enters the entity text “JNK-1”, then presses the “Ground with GILDA” button. (B) Gilda returns HGNC:6881 (standard name MAPK8) as the grounding, which is then used to perform the search.

Gilda is used to support human-machine dialogue in the context of the CLARE system which plugs into Slack workspaces as an application. In this context, Gilda grounds named entities appearing in questions from users, a key component of interpreting natural language input and constructing responses. Figure 3 shows an example dialogue in which Gilda is invoked to ground a human gene, a biological process, and a small molecule based on non-standard synonyms.

**Figure 3.**
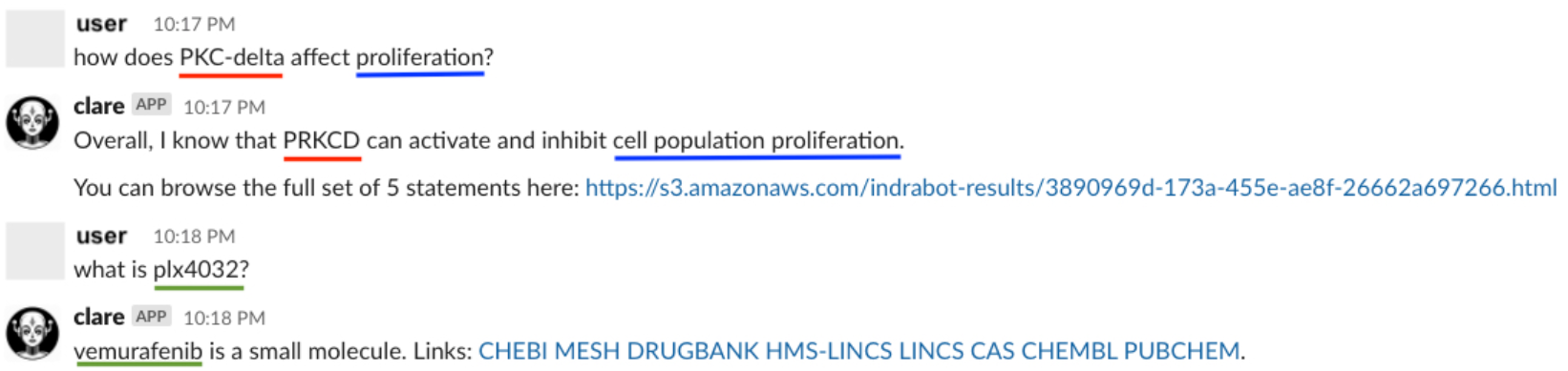
Gilda is used to ground three entity texts in the context of human-machine dialogue. “PKC-delta” is grounded to HGNC:9399 with standard name PRKCD (red underline), “proliferation” is grounded to GO:0008283 with standard name cell population proliferation (blue underline), and “plx4032” is grounded to CHEBI:63637 with standard name vemurafenib (green underline).

## Footnotes

1 https://github.com/pyobo/pyobo

2 https://biocreative.bioinformatics.udel.edu/tasks/biocreative-vi/track-1/

